# Genomic surveillance of antimicrobial-resistant *Escherichia coli* in fecal sludge and sewage in Uganda

**DOI:** 10.1101/2023.05.17.540885

**Authors:** Ryota Gomi, Yasufumi Matsumura, Masaki Yamamoto, Mai Tanaka, Allan John Komakech, Tomonari Matsuda, Hidenori Harada

## Abstract

The global increase of antimicrobial resistance (AMR) is a major public health concern. An effective AMR surveillance tool is needed to track the emergence and spread of AMR. Wastewater surveillance has been proposed as a resource-efficient tool for monitoring AMR carriage in the community. Here, we performed genomic surveillance of antimicrobial-resistant *Escherichia coli* obtained from fecal sludge and sewage in Uganda to gain insights into *E. coli* epidemiology and AMR burden in the underlying population. Selective media containing different antibiotic combinations (cefotaxime, ciprofloxacin, cefotaxime + ciprofloxacin + gentamicin) were used to obtain antimicrobial-resistant *E. coli* from fecal sludge and sewage. Short-read sequencing was performed for the obtained isolates, and a subset of isolates (selected from predominant sequence types (STs)) was also subjected to long-read sequencing. Genomic analysis of the obtained *E. coli* isolates (n = 181) revealed the prevalence of clonal complex 10, including ST167 (n = 43), ST10 (n = 28), ST1284 (n = 17), and ST617 (n = 4), in both fecal sludge and sewage, irrespective of antibiotics used for selection. We also detected global high-risk clones ST1193 (n = 10) and ST131 (n = 2 clade A, n = 3 subclade C1-M27, and n = 1 subclade C2). Diverse AMR determinants, including extended-spectrum β-lactamase genes (mostly *bla*_CTX-M-15_) and mutations in *gyrA* and *parC*, were identified. Analysis of the completed genomes revealed that diverse IncF plasmids and chromosomal integration were the major contributors to the spread of AMR genes in the predominant STs. This study showed that a combination of sewage surveillance (or fecal sludge surveillance) and whole-genome sequencing can be a powerful tool for monitoring AMR carriage in the underlying population.

## 1. Introduction

The increase of antimicrobial resistance (AMR) is a global health concern. AMR can lead to adverse consequences such as increased mortality rates, prolonged hospital admissions, and higher medical costs (Majumder et al., 2020). Information on AMR collected through surveillance is needed to guide treatment, monitor the effectiveness of interventions, and develop polices and regulations to counter AMR (WHO, 2015). However, surveillance of AMR in resource-limited settings is challenging, especially when the surveillance targets populations.

Sewage contains pooled urine and feces from a large population, and one sewage sample can represent bacteria from multiple people in the community. For this reason, sewage surveillance (or wastewater-based epidemiology) has been proposed as a resource-efficient tool for monitoring AMR carriage in the underlying population (Aarestrup and Woolhouse, 2020; Larsson et al., 2022). Sewage surveillance is attractive in many aspects; for example, (i) sewage provides a sample from a large and mostly healthy population, which otherwise would not be easily monitored, and (ii) it does not collect data on individuals and thus reduces ethical risks (Hendriksen et al., 2019). Although it is a promising approach, sewage surveillance of AMR cannot be directly applied in areas with no connection to centralized (sewered) systems. A previous study reported at least 1.8 billion people in low- and middle-income countries use on-site sanitation systems like pit latrines (Berendes et al., 2017). In such areas, fecal sludge from on-site systems has been proposed to be used as pooled community samples for surveillance purposes (Capone et al., 2020; Chigwechokha et al., 2022). Sewage surveillance (or fecal sludge surveillance) of AMR can be either culture-dependent or culture-independent. Culture-independent methods (e.g., short-read metagenomics) have some advantages over culture-dependent methods, such as the ability to detect thousands of AMR genes in a single sample. However, it provides limited information on the host or the genetic context of AMR genes (Hendriksen et al., 2019). Culture-dependent methods, on the other hand, can link AMR genes to specific host bacteria, provide information on phylogenetic characteristics on the host, and may even provide information on the location (chromosome or plasmid) and genetic context of AMR genes.

*Escherichia coli* is a commensal bacterium in humans and animals and is also known to cause serious infections (Denamur et al., 2021). Although *E. coli* is intrinsically susceptible to almost all clinically relevant antibiotics, it can accumulate AMR genes via, e.g., horizontal gene transfer (Poirel et al., 2018). *E. coli* has been suggested as an indicator for AMR surveillance in the environment (Anjum et al., 2021), and previous studies have shown that *E. coli* is a good target organism for sewage surveillance of AMR (Huijbers et al., 2020; Hutinel et al., 2019; Kwak et al., 2015; Raven et al., 2019). WHO also selected extended-spectrum β-lactamase (ESBL)-producing *E. coli* as an indicator for “One Health” AMR surveillance (including sewage surveillance) in the Tricycle protocol (WHO, 2021). The protocol recommends usage of whole-genome sequencing (if possible) to determine the sequence type (ST), virulence genotype, plasmid type, carriage of acquired AMR genes, and phylogenetic characteristics of the obtained isolates, which contribute to the understanding of the epidemiology of pathogens.

Uganda is a country experiencing an extremely high prevalence of AMR (Nabadda et al., 2021). Previous studies have reported high prevalence of multidrug resistance (MDR) among *E. coli* (Katongole et al., 2020; Odongo et al., 2020). However, most studies were based on phenotypic characterization or detection of a limited set of AMR genes/virulence genes, and thus their genomic characteristics, including the clonal composition and resistome, are largely unknown. In the present study, we isolated antimicrobial-resistant *E. coli* from fecal sludge and sewage in Uganda and performed genomic analysis on the obtained isolates, aiming to gain insights into *E. coli* epidemiology and AMR burden in the underlying population.

## 2. Materials and methods

### 2.1. Sample collection and isolation of *E. coli*

Samples were collected at a fecal sludge treatment plant (FSTP) and a wastewater treatment plant (WWTP) in Kampala, Uganda, in October–November, 2018. We obtained samples from both sources to cover a wider population in this region. For fecal sludge sampling, three grab samples were collected at different times on each of five days from effluent of the primary sedimentation tank in the FSTP; in total 15 samples were collected (these samples are hereafter referred to simply as fecal sludge samples). For sewage sampling, up to three grab samples were collected at different times over six days from influent of the WWTP, totaling 16 samples. The samples were collected and stored in sterile sampling bottles and transported on ice to the laboratory. The samples were then cultured, within 12 hours after collection, using a specific enzyme substrate medium (CHROMagar ECC, Kanto Chemical, Tokyo, Japan) containing four combinations of three antibiotics: (a) no antibiotics, (b) cefotaxime (CTX, 1 mg/L), (c) ciprofloxacin (CIP, 1 mg/L), (d) CTX (1 mg/L) + CIP (1 mg/L) + gentamicin (GM, 4 mg/L). The samples were serially diluted with PBS for colony counting. Up to three colonies with an *E. coli* profile (blue colony) were picked from each of plate (b), (c), and (d) (i.e., we obtained up to nine isolates from a single sample), and each was transferred into a screw tube containing LB agar. The tubes were incubated overnight at 37 °C, stored at 4 °C, and then transported to the laboratory in Japan. *E. coli* in each screw tube was again incubated on a plate with CHROMAgar ECC containing the same antibiotic(s) as at the time of isolation. A colony with an *E. coli* profile was picked and spread again on a plate with CHROMAgar ECC, and then a single colony was picked from the plate, cultured in LB broth, and stored at –85 °C in 35% glycerol.

### 2.2. Genome sequencing and assembly

A total of 192 presumptive antimicrobial-resistant *E. coli* isolates was subjected to genome sequencing as below. DNA was extracted from each isolate using a QIAamp DNA Mini Kit (Qiagen, Hilden, Germany) for Illumina sequencing. Illumina libraries were prepared using a Nextera XT DNA Library Preparation Kit (Illumina, San Diego, CA) and sequenced on the NextSeq550 platform, generating paired-end reads of 151 bp each. For some isolates, read depth was not sufficient, so we sequenced the library again on the MiniSeq platform to obtain additional paired-end reads of the same length. Illumina reads were processed using fastp (v0.23.2) (Chen et al., 2018) and assembled using Unicycler (v0.4.8) with the --no_correct option (Wick et al., 2017). The quality of each genome assembly was ensured by using the quality control (QC) criteria of (i) contig number: ≤ 800; (ii) genome size: 3.7 Mbp to 6.4 Mbp; and (iii) N50: >20 kb, as recommended by EnteroBase (Zhou et al., 2020).

For selected isolates (n = 26), we also performed Oxford Nanopore Technologies (ONT) long-read sequencing to generate complete genomes. DNA was extracted from each isolate using a DNeasy PowerSoil Pro Kit (Qiagen). The long-read sequencing library was prepared using the SQK-LSK109 kit (Oxford Nanopore Technologies, Oxford, UK) and sequenced on the MinION with a FLO-MIN106 flow cell. ONT reads were filtered using Filtlong (v0.2.1, https://github.com/rrwick/Filtlong) to remove reads with low quality or short length. Hybrid assembly of Illumina short reads and ONT long reads was performed by employing a long-read-first approach. Briefly, filtered ONT reads were assembled using Flye (v2.9.1-b1780) (Kolmogorov et al., 2019), and then the assemblies were polished with medaka (v1.7.1, https://github.com/nanoporetech/medaka) using filtered ONT reads and Polypolish (v0.5.0) using processed Illumina reads (Wick and Holt, 2022). Hybrid assembly was also performed using a short-read-first approach, i.e., Unicycler (v0.5.0) hybrid assembly followed by polishing with Polypolish (v0.5.0) using processed Illumina reads. If the long-read-first approach generated a complete genome for an isolate, we chose the polished Flye assembly as the final assembly. If the long-read-first approach did not complete the genome but the short-read-first approach did, we chose the polished Unicycler assembly as the final assembly.

### 2.3. Genomic analysis

*E. coli* status was confirmed by performing an average nucleotide identity (ANI) analysis using each genome sequenced in the present study and the genome sequence of the *E. coli* type strain ATCC 11775 (= DSM 30083 = U5/41) (Wadley et al., 2019). ANI analysis was performed using FastANI (v1.33) with the threshold of 95% (Jain et al., 2018). *E. coli* phylogroups were determined using a combination of EzClermont (v0.7.0) and ClermonTyping (Beghain et al., 2018; Waters et al., 2020). Multilocus sequence typing (MLST) was performed using MLST (v2.19.0) (https://github.com/tseemann/mlst). *fimH* types, which are used for high-resolution subtyping of MLST-based *E. coli* clonal groups, were determined using FimTyper 1.0 (Roer et al., 2017). AMR genes were detected using a combination of ResFinder 4.1 (Bortolaia et al., 2020) and Abricate (v1.0.1) (https://github.com/tseemann/abricate) with the ncbi database. Genetic contexts of AMR genes were analyzed using the RAST server (Aziz et al., 2008), the BLAST algorithm (https://blast.ncbi.nlm.nih.gov/Blast.cgi), and ISfinder (Siguier et al., 2006). Mutations in quinolone resistance-determining regions (QRDRs) in *gyrA* (GyrA codons 83 and 87) and *parC* (ParC codons 80 and 84) were analyzed as described previously (Aoike et al., 2013). Virulence genes were detected using the VirulenceFinder database (Joensen et al., 2014; Malberg Tetzschner et al., 2020). *E. coli* pathotypes, which include extraintestinal pathogenic *E. coli* (ExPEC) and several categories of intestinal pathogenic *E. coli* (InPEC), were defined based on the presence of specific virulence genes as described previously (Gomi et al., 2015). K1 and K5 capsular types of ST1193 isolates were determined based on blastn analysis using M57382 (K1) and X53819 (K5) as references (Johnson and Stell, 2000). Plasmid replicons were detected using PlasmidFinder 2.1, and plasmids were typed using pMLST 2.0 (Carattoli et al., 2014).

A whole-genome SNP-based tree was built for isolates belonging to phylogroup A using kSNP3 based on SNPs occurring in at least 90% of the genomes (Gardner et al., 2015). To confirm the topology of the tree, we also built a tree for the phylogroup A genomes using another software, parsnp (v1.7.4) (Treangen et al., 2014). For isolates belonging to clonal complex 10 (CC10) within phylogroup A, we built a tree together with global CC10 genomes (n = 4,744) obtained from the NCBI Reference Sequence (RefSeq) database. For isolates belonging to ST1193 and ST131, we separately built a recombination-free phylogenetic tree for each ST. To place our isolates in broader phylogenetic contexts, we also included isolates analyzed in other studies which defined clades/subclades within each of ST1193 and ST131 (Johnson et al., 2019; Matsumura et al., 2017b). Detailed methods used for tree construction of the CC10, ST1193, and ST131 genomes are provided in the Supplementary Materials and methods.

### 2.4. Accession numbers

Genome assemblies were deposited in GenBank under BioProject PRJNA956323 (also see **Table S1** for the accession number of each genome and **Table S2** for the accession number of each replicon for the completed genomes).

## 3. Results and discussion

### 3.1. Detection and isolation of antimicrobial-resistant *E. coli*

During the study period, we collected 15 fecal sludge samples and 16 sewage samples and measured the concentrations of total *E. coli* and antimicrobial-resistant *E. coli* (**Table S3**). The mean proportions of antimicrobial-resistant *E. coli* in the fecal sludge samples were 30.0% for CTX, 26.9% for CIP, and 10.7% for CTX + CIP + GM. The mean proportions of antimicrobial-resistant *E. coli* in the sewage samples were 39.2% for CTX, 21.0% for CIP, and 5.1% for CTX + CIP + GM. These proportions were higher than those reported previously for *E. coli* in wastewater influent in some developed countries; for example, the proportions of antimicrobial-resistant *E. coli* were 0%–3.2% for CTX and 3.2%–7.1% for CIP in Sweden (Hutinel et al., 2019) and 1.6%–4.4% for CTX (or ceftazidime) and 2.4%–13.4% for CIP in ten European countries (Huijbers et al., 2020). On the other hand, the proportions were comparable to those reported in some other low- and middle-income countries (LMICs); for example, the resistance proportions were 38% for CTX and 35% for CIP in South Africa (Gumede et al., 2021) and 24% for CIP in Vietnam (Le et al., 2023), potentially highlighting the extremely high prevalence of AMR in Uganda and some LMICs (Nabadda et al., 2021; WHO, 2022).

In total, we obtained 192 presumptive antimicrobial-resistant *E. coli* isolates from the samples and performed short-read sequencing on these isolates. After discarding contigs shorter than 100 bp, 187 genomes met the assembly QC criteria. Species identification of the 187 genomes by the ANI analysis revealed that 181 belonged to *E. coli*, while 6 belonged to non-*E. coli* species such as *Citrobacter freundii*. These 181 *E. coli* genomes, comprising 91 from fecal sludge samples (31 from CTX-containing media, 31 from CIP-containing media, and 29 from CTX + CIP + GM-containing media) and 90 from sewage samples (32 from CTX-containing media, 26 from CIP-containing media, and 32 from CTX + CIP + GM-containing media), were further analyzed in the present study.

### 3.2. Clonal composition

In total, 36 STs from six different phylogroups were detected among the 181 isolates (**Table S1**). Phylogroup A was the most prevalent (n = 127), followed by B1 (n = 17), B2 (n = 17), C (n = 10), D (n = 8), and F (n = 2). ST167 (n = 43), ST10 (n = 28), ST46 (n = 19), and ST1284 (n = 17), all of which belong to phylogroup A, were prevalent in both fecal sludge isolates and sewage isolates (**Figure 1a**). In total, 144 isolates (79.6%) were from the top ten STs, while 37 isolates (20.4%) were from minor STs (i.e., STs that are not in the top ten and detected among three or fewer isolates) (**Figures 1b** and **1c**). The top ten STs were generally associated with two or three types of media and both fecal sludge and sewage, with a few exceptions (e.g., ST131 was exclusively found among fecal sludge isolates). Lineages associated with extraintestinal infections were also among the top ten STs and detected in fecal sludge isolates (n = 9 ST1193, n = 5 ST410, n = 6 ST131, n = 2 ST69) and sewage isolates (n = 1 ST1193, n = 4 ST410, n = 2 ST69) (Denamur et al., 2021). Thirty-four isolates (18.8%), including those from the above-mentioned STs associated with extraintestinal infections, were classified as ExPEC based on the virulence gene contents (**Figure 1**). On the other hand, none of the isolates were classified as InPEC. ExPEC strains are facultative pathogens and part of the intestinal microflora of a fraction of the healthy population (Kohler and Dobrindt, 2011), which might explain the relatively high prevalence of ExPEC among our *E. coli* collection.

**Figure 1.**
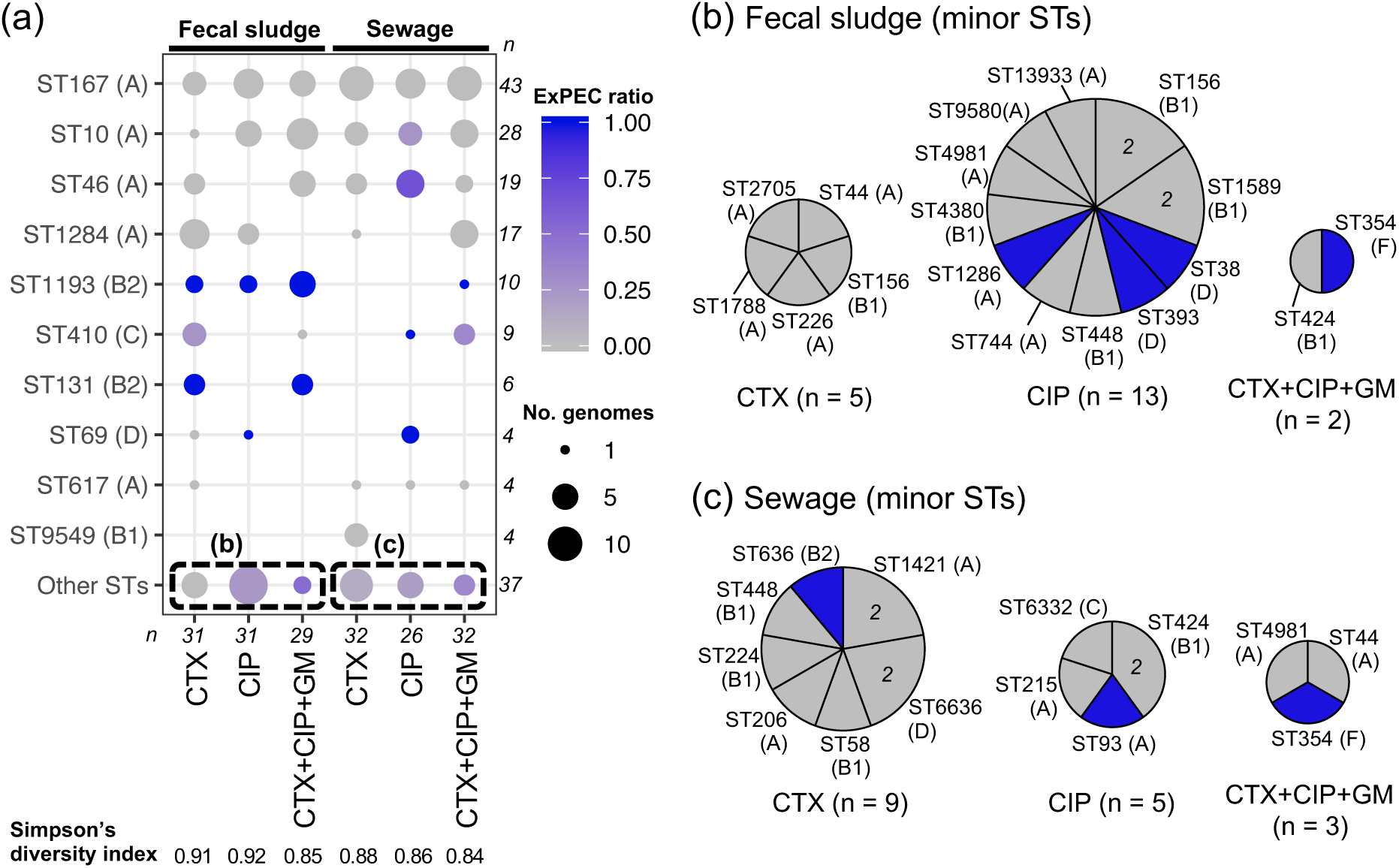
(a) Clonal composition of 181 *E. coli* isolates. STs are shown separately for each media type (CTX, CIP, and CTX + CIP + GM) and each sample type (fecal sludge and sewage). The top ten STs are shown in descending order. The remaining STs, all of which were with three or fewer isolates, are combined into “other STs”. Phylogroup corresponding to each ST is shown in parentheses. Circle size indicates the number of genomes, and intensity of circle shading indicates the proportion of genomes classified as ExPEC (as per inset legends). Clonal composition of minor STs (i.e., STs classified as “other STs” in (a)) from (b) fecal sludge and (c) sewage. Phylogroup corresponding to each ST is shown in parentheses, and STs classified as ExPEC are colored blue.

We next compared the clonal composition of *E. coli* in this study and those reported previously for clinical *E. coli* in Uganda. However, the clonal composition of *E. coli* in clinical settings is largely unknown in Uganda (a search of human-derived *E. coli* genomes from Uganda in EnteroBase returned no hits as of May 2023), and we could find only one study reporting the ST composition of clinical *E. coli* isolates in Uganda. That study sequenced 24 Ugandan *E. coli* isolates causing urinary tract infections (most were MDR) and reported the prevalence of ST10 (n = 4, 15.4%) as well as the occurrence of ST410 (n = 2, 8.3%), ST131 (n = 1, 4.2%), ST167 (n = 1, 4.2%), ST617 (n = 1, 4.2%), and ST1193 (n = 1, 4.2%) (Decano et al., 2021). These STs were also among the top ten STs in the present study, indicating that sewage surveillance may serve as a possible resource-efficient complement to clinical surveillance, though further studies are needed to confirm this.

Previous studies have suggested that fecal sludge and sewage can be used for community surveillance, but fecal sludge includes waste from fewer individuals and hence may be less divergent than sewage (Capone et al., 2020). However, the number of unique STs was larger for fecal sludge isolates than sewage isolates (26 unique STs vs 23 unique STs) (**Table S1**). Moreover, Simpson’s diversity index was higher for fecal sludge isolates than sewage isolates when isolates from the same type of media were compared (**Figure 1a**). This might be because of the fecal sludge sampling strategy employed in the present study: we obtained samples from effluent of the primary sedimentation tank in a FSTP, which probably represents fecal sludge from a large number of individuals. This might have contributed to the relatively high diversity of *E. coli* isolates from fecal sludge samples, suggesting sampling from FSTPs or otherwise designated fecal sludge dumping sites (rather than individual pit latrines) may better capture the population-scale information in fecal sludge surveillance, when such places exist in the region.

### 3.3. Antimicrobial resistance determinants

Six different ESBL genes, namely *bla*_CTX-M-3_, *bla*_CTX-M-14_, *bla*_CTX-M-15_, *bla*_CTX-M-27_, *bla*_CTX-M-55_, and *bla*_TEM-169_, were detected among the isolates, with *bla*_CTX-M-15_ being the predominant type (**Figure 2a**). Based on the short-read assemblies, there were 21 isolates with two different ESBL genes (n = 18 *bla*_CTX-M-14_ + *bla*_CTX-M-15_, n = 1 *bla*_CTX-M-15_ + *bla*_CTX-M-27_, n = 1 *bla*_CTX-M-3_ + *bla*_CTX-M-27_, and n = 1 *bla*_TEM-169_ + *bla*_CTX-M-15_) (note that isolates carrying multiple copies of the same ESBL gene were not considered here because those genes can be collapsed into one contig in short-read assemblies). As expected, all but one isolate obtained from CTX-containing media and CTX + CIP + GM-containing media carried at least one ESBL gene. Importantly, many isolates obtained from CIP-containing media carried ESBL genes (41.9% for fecal sludge isolates and 42.3% for sewage isolates), indicating the convergence of ESBL genes and fluoroquinolone resistance.

**Figure 2.**
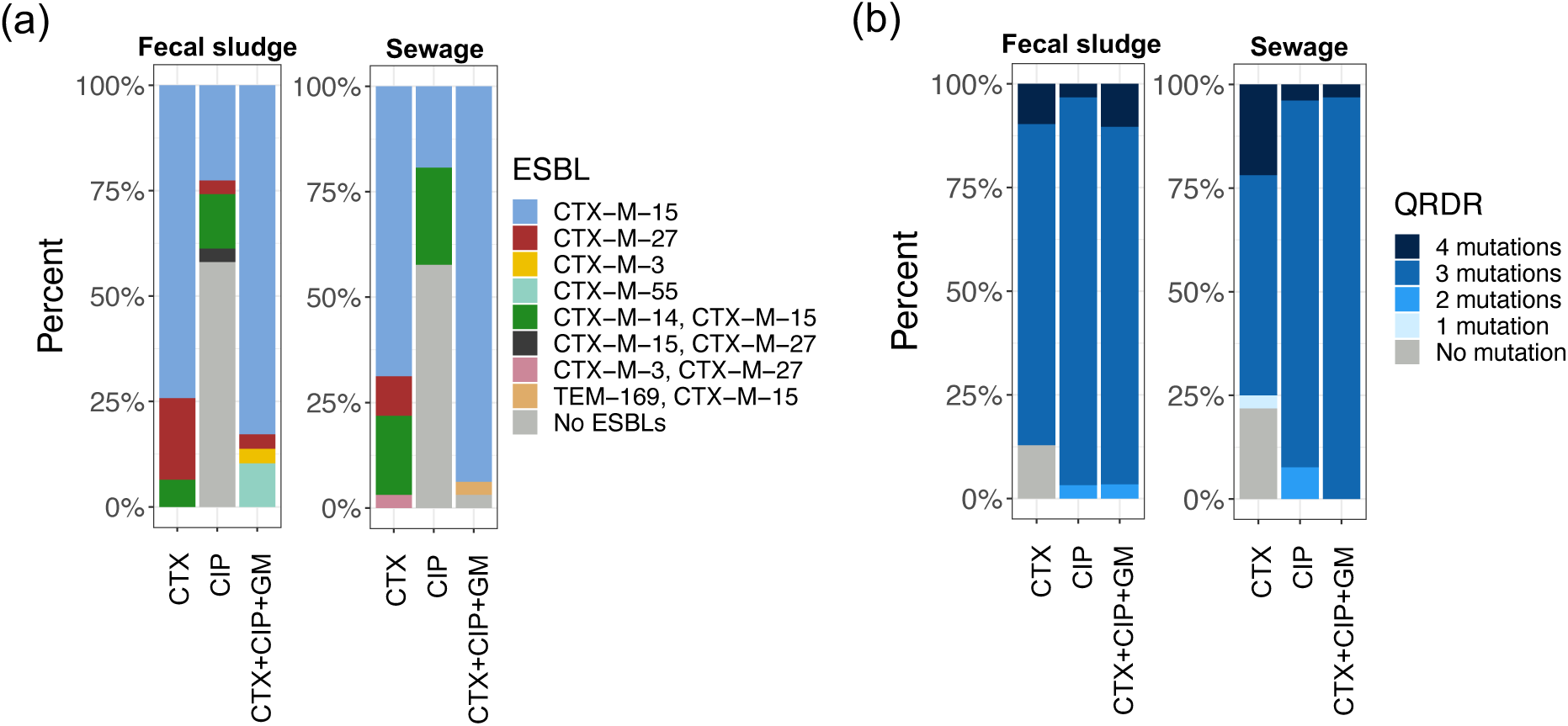
(a) ESBL genes and (b) QRDR mutations in *E. coli* isolates. AMR determinants are shown separately for each media type (CTX, CIP, and CTX + CIP + GM) and each sample type (fecal sludge and sewage). This figure is based on the analysis of short-read assembly data and isolates with multiple copies of the same ESBL gene can be reported to carry a single copy. For QRDRs, mutations in GyrA codons 83 and 87 and ParC codons 80 and 84 were counted.

All but four isolates obtained from CIP-containing media and CTX + CIP + GM-containing media carried three or four mutations in QRDRs (**Figure 2b**). In all isolates with three mutations in QRDRs, two were in codons 83 and 87 in GyrA, and the remaining mutation was in ParC codon 80 or 84. These are congruent with previous studies reporting that two mutations in the QRDR of *gyrA* and one or two mutations in the QRDR of *parC* are the main mechanism of ciprofloxacin resistance in *E. coli* (Matsumura et al., 2017a; van der Putten et al., 2019). Importantly, most isolates obtained from CTX-containing media also carried three or four mutations in QRDRs (87.1% for fecal sludge isolates and 75.0% for sewage isolates), again highlighting the convergence of ESBL genes and fluoroquinolone resistance determinants.

Gentamicin resistance in isolates obtained from CTX + CIP + GM-containing media could be mostly explained by the presence of *aac(3)-II* genes (**Table S1**). Other resistance genes not associated with antibiotics used in the selective media were also prevalent in our isolates; for example, *sul* genes were detected in 175 isolates (96.7%) and *tet* genes were detected in 161 isolates (89.0%). However, genes conferring resistance to last-resort antibiotics, such as carbapenems and colistin, were not detected in any isolates (**Table S1**). We did not use last-resort antibiotics to select *E. coli* isolates in the present study, but we note that large-scale genomic surveillance of isolates resistant to last-resort antibiotics should also be performed in LMICs in future studies.

### 3.4. Genomic characteristics of phylogroup A isolates

Whole genome-based phylogenetic analysis was performed using kSNP3 to gain insights into clonal relationships of 127 isolates belonging to phylogroup A, which was the dominant phylogroup in the present study (**Figure 3**). The tree was largely separated according to STs, though we observed separate clusters for ST10, ST46, and ST167. These separate clusters were also confirmed in a tree generated using different software, parsnp (**Figure S1**). The ST10 and ST167 clusters were characterized by distinct *fimH* types, namely *fimH54*, *fimH435*, and *fimH789* for ST10, and *fimH5*, *fimH29*, and *fimH54* for ST167. ST46 isolates from both clusters did not carry *fimH*.

**Figure 3.**
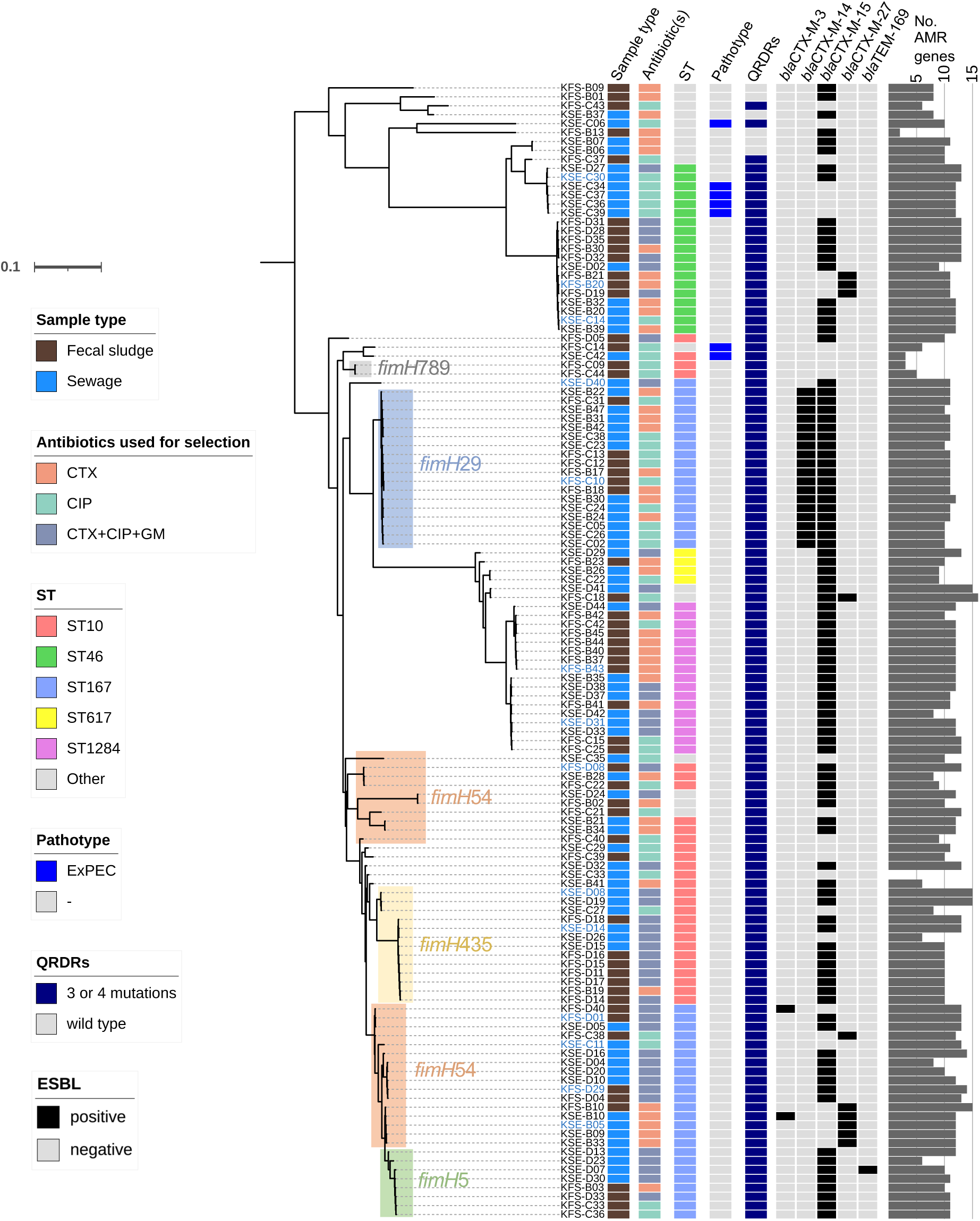
kSNP3 whole-genome SNP-based phylogenetic tree of 127 phylogroup A isolates. The tree was constructed using 54,632 SNP-loci, which occurred in at least 90% of the genomes. A k-mer length of 19 was used for SNP identification. The presence of traits is indicated by colored boxes (as per inset legends). For the ST column, major STs (i.e., ST10, ST46, ST167, ST617, and ST1284) are indicated by the corresponding color boxes, while other STs are represented by gray boxes. ST10 and ST167 clusters are highlighted in the tree according to the *fimH* alleles, except for singletons. The names of completed genomes are colored blue.

According to EnteroBase (Zhou et al., 2020), ST167, ST10, ST1284, and ST617 are all classified into CC10. Classifying all isolates from a monophyletic clade containing these four STs (the bottom clade in **Figure 3**) into CC10 identified that 99 isolates (54.7% of isolates in the present study) belong to this lineage, indicating CC10 poses a significant AMR burden in the underlying population. CC10 has been globally reported as a resident of the intestinal tract of humans and other animals, and carriage of AMR genes (including ESBL genes) has also been widely reported in this lineage (Reid et al., 2019). To determine the relationships between CC10 isolates in this study and global CC10 isolates, we built a Mash-based tree using our CC10 genomes and RefSeq CC10 genomes (**Figure S2** and **Table S4**). We found that our CC10 genomes formed clusters with global CC10 genomes (e.g., ST617-ST1284 cluster and ST167 cluster in **Figure S2**). On the other hand, some of the top 15 STs detected in global CC10 genomes (e.g., ST48 and ST746) were not detected in our CC10 genomes. We also found that our CC10 genomes were not evenly distributed in the tree, while African RefSeq CC10 genomes were scattered throughout the tree. These findings indicate that only a subset of global clones within CC10 were prevalent in our study setting.

Although there were some small clusters specific to fecal sludge isolates (e.g., ST10 with *fimH789*) and sewage isolates (e.g., a cluster containing six sewage isolates within ST46), fecal sludge isolates and sewage isolates were generally intermingled in the tree (**Figure 3**). This trend was also seen for media types; isolates selected with different antibiotics were generally intermingled in the tree. Most phylogroup A isolates (except six isolates from minor STs obtained using CTX-containing media) carried three or four mutations within QRDRs. As mentioned above, ESBL genes were also prevalent even in isolates obtained using CIP-containing media (n = 21/40, 52.5%). Notably, all ST167 isolates with *fimH*29 (n = 18) carried both *bla*_CTX-M-14_ and *bla*_CTX-M-15_. Analysis of the completed genome of KFS-C10 from this clade identified that *bla*_CTX-M-14_ was integrated into the chromosome while *bla*_CTX-M-15_ was carried on an IncF[F31:F36:A4:B1] plasmid (note that this plasmid carried two different FII alleles, F31 and F36) (**Table S2**). There were seven isolates with ExPEC status (n = 4 ST46, n = 1 ST10, n = 1 ST93, n = 1 ST1286). All seven isolates were obtained from CIP-containing media and lacked ESBL genes, but they carried 3 to 12 non-ESBL AMR genes in addition to QRDR mutations.

### 3.5. Genomic characteristics of ST1193 and ST131 isolates

We investigated the detailed genetic characteristics of isolates belonging to ST1193 (n = 10) and ST131 (n = 6), most of which were obtained from fecal sludge, because these STs are recognized as established/emerging global high-risk clones and the only MDR clones that are dominant among unselected ExPEC populations (Pitout et al., 2022).

ST1193 is derived from the ST14 clonal complex, carries *fimH64*, and harbors *gyrA* (S83L and D87N) and *parC* (S80I) mutations (Pitout et al., 2022). Previous studies reported that ST1193 isolates cluster together within the phylogenetic tree according to three major IncF plasmid types, namely F-:A1:B1, F-:A1:B10, and F-:A1:B20, though the clusters are not strictly monophyletic (Johnson et al., 2019; Wyrsch et al., 2022). These clusters were named as green (F-:A1:B1), red (F-:A1:B10), and pink (F-:A1:B20) (Johnson et al., 2019). The phylogenetic tree constructed in our study was congruent with these previous studies (i.e., largely separated according to IncF types), and our isolates fell into an F-:A1:B10 cluster (n = 7) and an F-:A1:B20 cluster (n = 3) (**Figure 4a**). Three isolates (KFS-B24, KFS-B25, KFS-C19) and two isolates (KFS-D07, KSE-D22) with the F-:A1:B10 plasmid type clustered together in the tree. Three isolates (KFS-D21, KFS-D24, KFS-D26) with the F-:A1:B20 plasmid type also clustered together. ST1193 isolates with F-:A1:B:20 plasmids generally contain the K5 capsular type, and those with F-:A1:B1 or F-:A1:B10 plasmids generally contain the K1 capsular type (Johnson et al., 2019). However, all of our ST1193 isolates, including those with F-:A1:B:20 plasmids, contained the K1 capsular type. A variety of AMR genes associated with different drug classes were detected in our ST1193 isolates (**Figure 4c**). Notably, among six isolates with complete genomes, three (KFS-B24, KFS-D13, KSE-D22) carried chromosomal AMR genes, including *bla*_CTX-M-15_, within different genetic contexts (discussed in detail below and in **Figure S3**). To determine if global ST1193 genomes carry chromosomal AMR genes, we checked the location of AMR genes in all complete ST1193 genomes deposited in GenBank (as of October 2023). Among 46 genomes, excluding our own genomes, eight (17%) carried chromosomal AMR genes and six (13%) carried chromosomal *bla*_CTX-M_ (**Table S5**). We also found that the genetic contexts of chromosomal *bla*_CTX-M_ in ST1193 genomes were diverse and the contexts detected in two of our ST1193 genomes (KFS-B24 and KFS-D13) were not described previously, indicating multiple events contributed to integration of *bla*_CTX-M_ into the ST1193 chromosome. The transposition unit detected in the chromosome of KSE-D22, which contains *bla*_CTX-M-15_ and four other AMR genes, was found to be inserted into the same position in the chromosome of a Canadian ST1193 *E. coli* genome (GCA_032296485.1), potentially suggesting global circulation of this strain. These observations indicate chromosomal integration of AMR genes, as well as plasmid carriage, can contribute to the MDR phenotype of ST1193. All ST1193 isolates in this study were classified as ExPEC based on the virulence gene profiles, which potentially indicates these isolates can cause extraintestinal infections.

**Figure 4.**
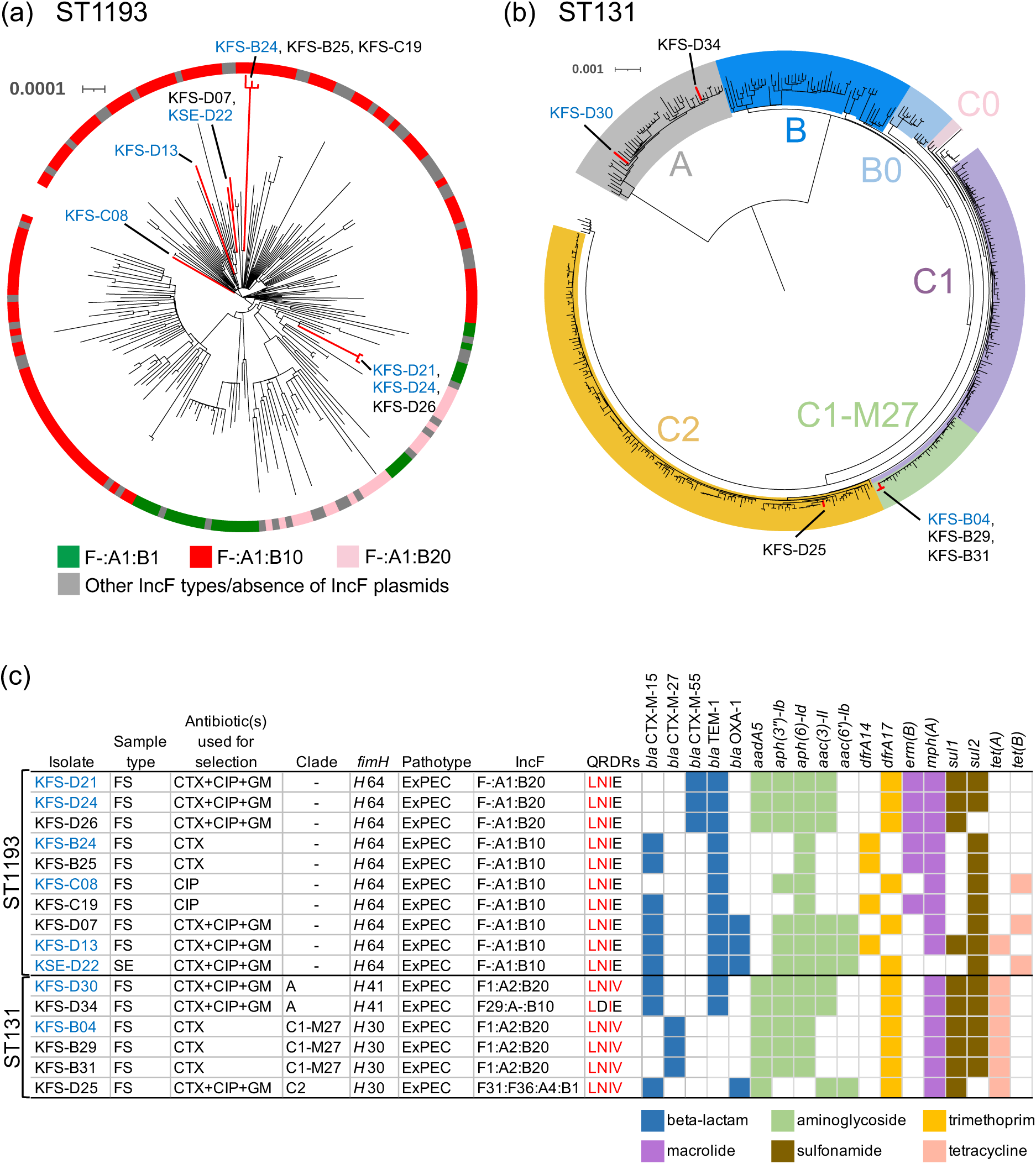
Recombination-free phylogenetic trees for (a) ST1193 and (b) ST131. For ST1193, we used the Assembly Dereplicator tool (v0.3.0, https://github.com/rrwick/Assembly-Dereplicator) and subsampled global isolates to remove near-identical redundant isolates and to reduce computational burden. The ST1193 tree was built using a 3,899 bp alignment generated from our isolates (n = 10) and global isolates (n = 198). The ST131 tree was built using a 10,712 bp alignment generated from our isolates (n = 6) and global isolates (n = 379). Branches leading to our isolates are colored in red. The trees were midpoint rooted. Accession numbers of global ST1193 genomes and ST131 genomes are available in **Table S6** and **Table S7**. (c) Key genetic features of ST1193 and ST131 isolates sequenced in the present study. FS, fecal sludge; SE, sewage. We did not assign clades to ST1193 isolates because the previously defined ST1193 clades are not monophyletic as discussed in the main text. For QRDRs, GyrA codons 83 and 87 and ParC codons 80 and 84 that are different from the wild type (SDSE) are colored in red. Presence of AMR genes is indicated by colored boxes (colored by drug class). All four isolates with *aac(6′)-Ib* carried the *aac(6′)-Ib-cr* variant, which is associated with decreased susceptibility to aminoglycosides and to ciprofloxacin and norfloxacin. KFS-D13 carried *aadA1*, *dfrA5*, *ere(A)*, *qnrB1*, and *catA1* in addition to AMR genes shown in (c). Genomes with blue names in (a)–(c) were completed.

ST131 is known to be divided into clades A, B, and C (containing subclades C0, C1, and C2) (Pitout and Finn, 2020). Within subclade C1 there is a clone known as C1-M27, which is responsible for the spread of ESBL-producing *E. coli* in some countries (Matsumura et al., 2017b; Merino et al., 2018). The ST131 isolates in the present study belonged to subclade C1-M27 (n = 3), clade A (n = 2), and subclade C2 (n = 1) (**Figure 4b**). Three subclade C1-M27 isolates clustered together within the tree, while two clade A isolates were distantly related. As reported previously (Matsumura et al., 2017b), the C1-M27 isolates carried *fimH*30, LNIV-type QRDRs, and *bla*_CTX-M-27_ (**Figure 4c**). The two clade A isolates were typed as *fimH41* as previously reported (Matsumura et al., 2017b), but carried *bla*_CTX-M-15_ and mutations in both *gyrA* and *parC* QRDRs. Although clade A *E. coli* is generally sensitive to antibiotics (Pitout and Finn, 2020), clade A isolates with *bla*_CTX-M_ and *gyrA*/*parC* mutations have been sporadically reported (Stoesser et al., 2016). The subclade C2 isolate carried *fimH*30, LNIV-type QRDRs, and *bla*_CTX-M-15_, which are the expected traits of the subclade C2 *E. coli* (Pitout and Finn, 2020). A variety of AMR genes were detected in our ST131 isolates, and all the isolates met the ExPEC criteria (**Figure 4c**).

Previous studies also identified the high-risk clones ST1193 and ST131 in sewage (Gomi et al., 2017; Raven et al., 2019), indicating sewage surveillance can be an effective tool to monitor the occurrence of these high-risk pathogens among populations. Genomic analysis, as implemented in the present study, resolved fine-scale phylogenies of these high-risk clones at the clade/subclade level (**Figure 4**), showing a combination of sewage (fecal sludge) surveillance and whole-genome sequencing is even more effective and informative.

### 3.6. Locations and genetic contexts of AMR genes in isolates from major clones

In total, 26 genomes were selected from the top five STs (n = 8 ST167, n = 4 ST10, n = 3 ST46, n = 2 ST1284, and n = 6 ST1193) and ST131 (n = 3) and subjected to long-read sequencing to elucidate the locations and genetic contexts of the AMR genes in these major STs (care was taken not to select multiple closely-related isolates). Hybrid assembly using short and long reads completed 22 of 26 genomes (n = 6 ST167, n = 3 ST10, n = 3 ST46, n = 2 ST1284, n = 6 ST1193, and n = 2 ST131). The remaining four genomes could not be completed, probably due to the presence of long repeats in the genomes.

The completed genomes carried one to eight plasmids (median four plasmids). Among the 89 plasmids in the completed genomes, 38 (42.7%) carried at least one AMR gene and thus were defined as AMR plasmids. IncF AMR plasmids were the most prevalent (n = 25), followed by Col(pHAD28) (n = 6), IncB/O/K/Z (n = 4), IncX1 (n = 1), IncX2 (n = 1), and IncHI2-IncHI2A (n = 1) (**Table 1**). We note that the completed genomes represent only a subset of our isolates, and the actual AMR plasmid diversity in the whole population may be different. Nonetheless, the prevalence of AMR IncF plasmids is noteworthy, and 11 (79%) of 14 plasmidic *bla*_CTX-M_ genes were located on IncF plasmids. IncF plasmids are frequently detected among *E. coli* from humans and can stably be maintained in commensal *E. coli* within the human gastrointestinal tract without antibiotic selection pressure (Bevan et al., 2017), which might explain this high prevalence. IncF replicon sequence typing identified 14 different allele combinations, revealing diverse IncF plasmids contributed to the spread of AMR genes within these major STs. Although AMR plasmids of other replicon types were not common, two IncB/O/K/Z plasmids and one IncHI2-IncHI2A plasmid contained a *bla*_CTX-M_ gene. We found plasmids from other continents that were highly similar to 13 (34%) of 38 AMR plasmids in our study under the stringent criteria applied (i.e., >99% query coverage and >99 % identity) (**Table 1**, **Table S2**). This indicates circulation of global AMR plasmids (or clones with such plasmids) in our study setting. On the other hand, close blastn hits (defined here as hits with >80% query coverage) were not found for some AMR plasmids (CP124985, CP124987, and CP124993), indicating the occurrence of unique/rare AMR plasmids in this region.

**Table 1.**
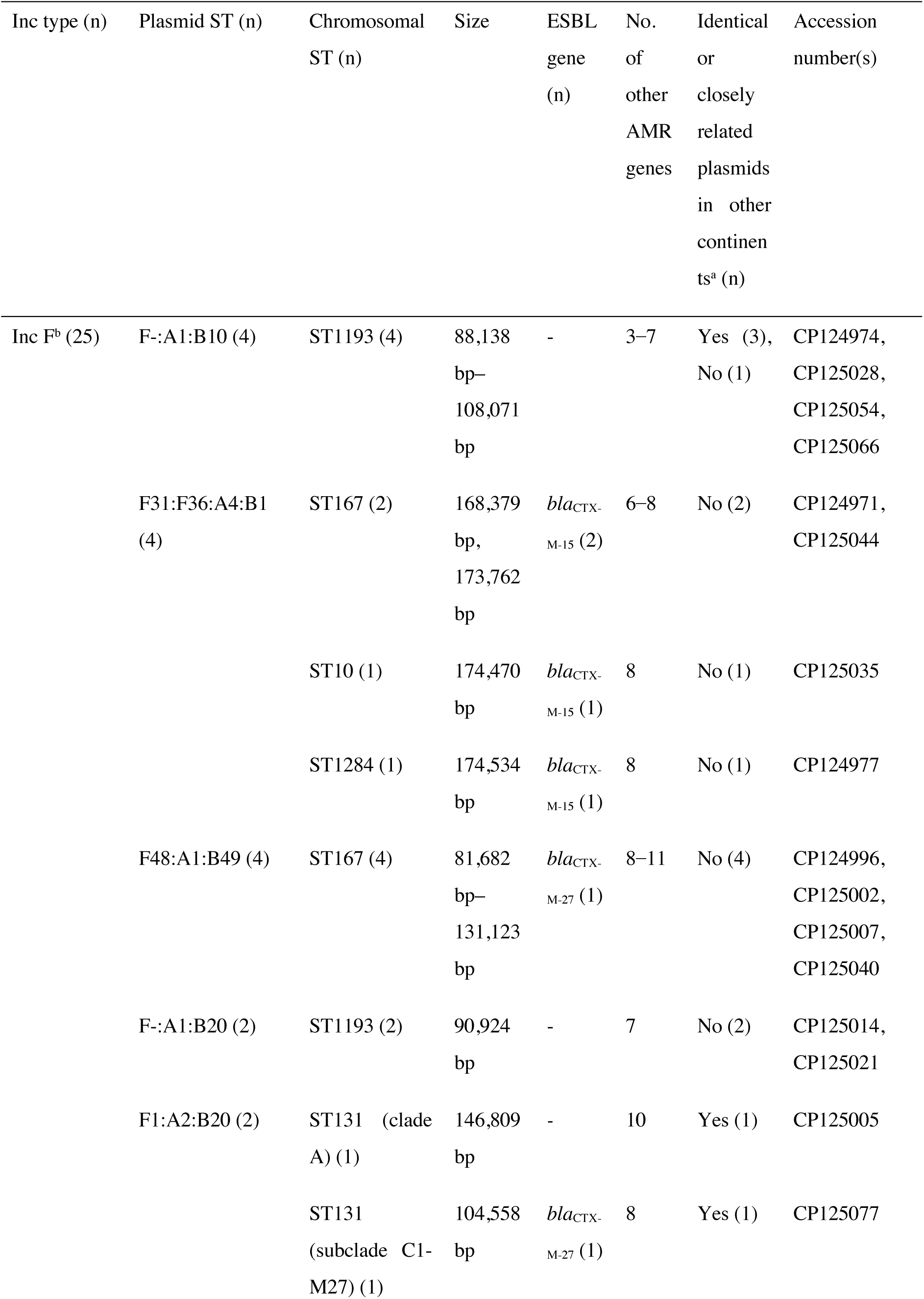
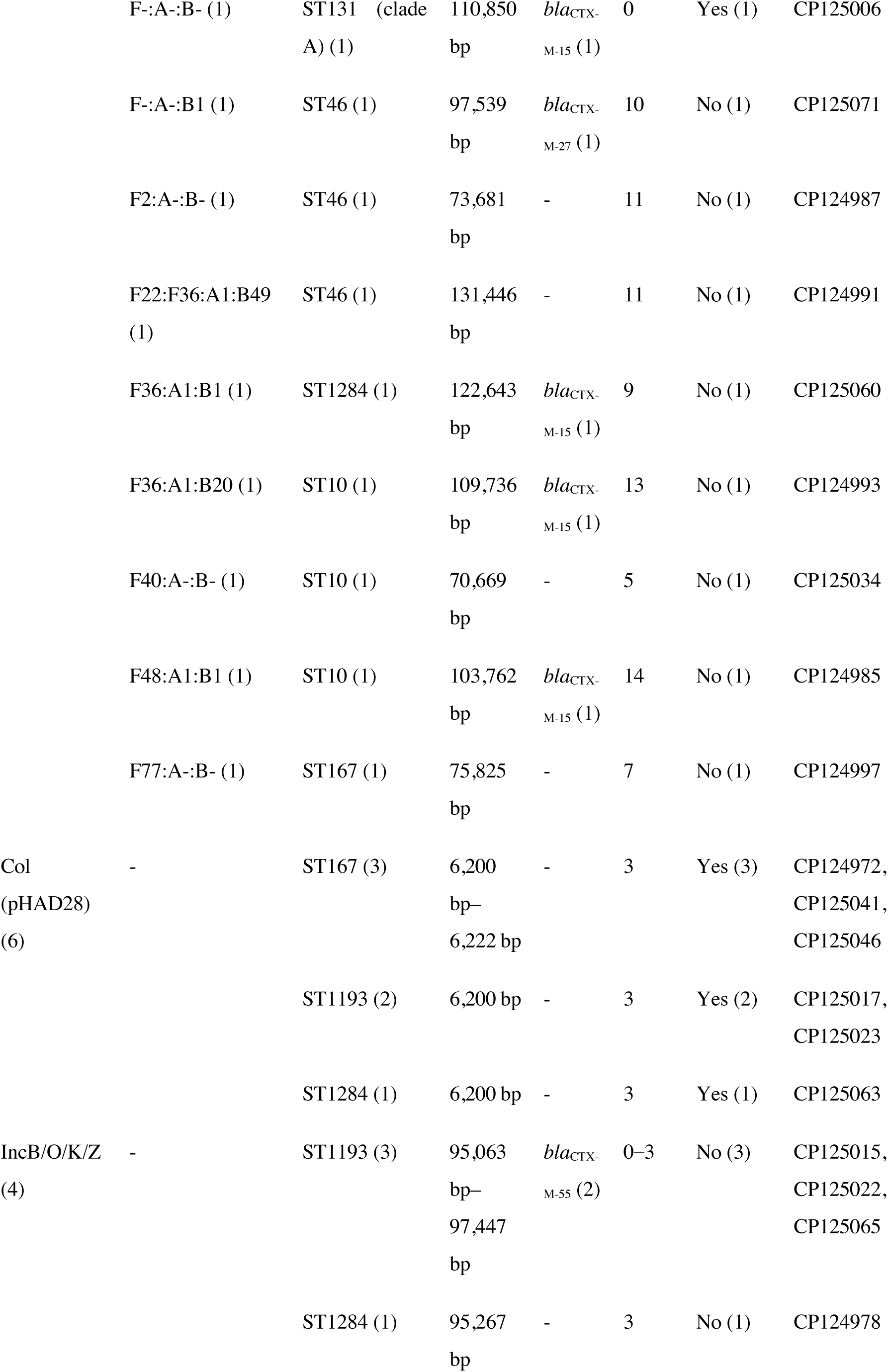
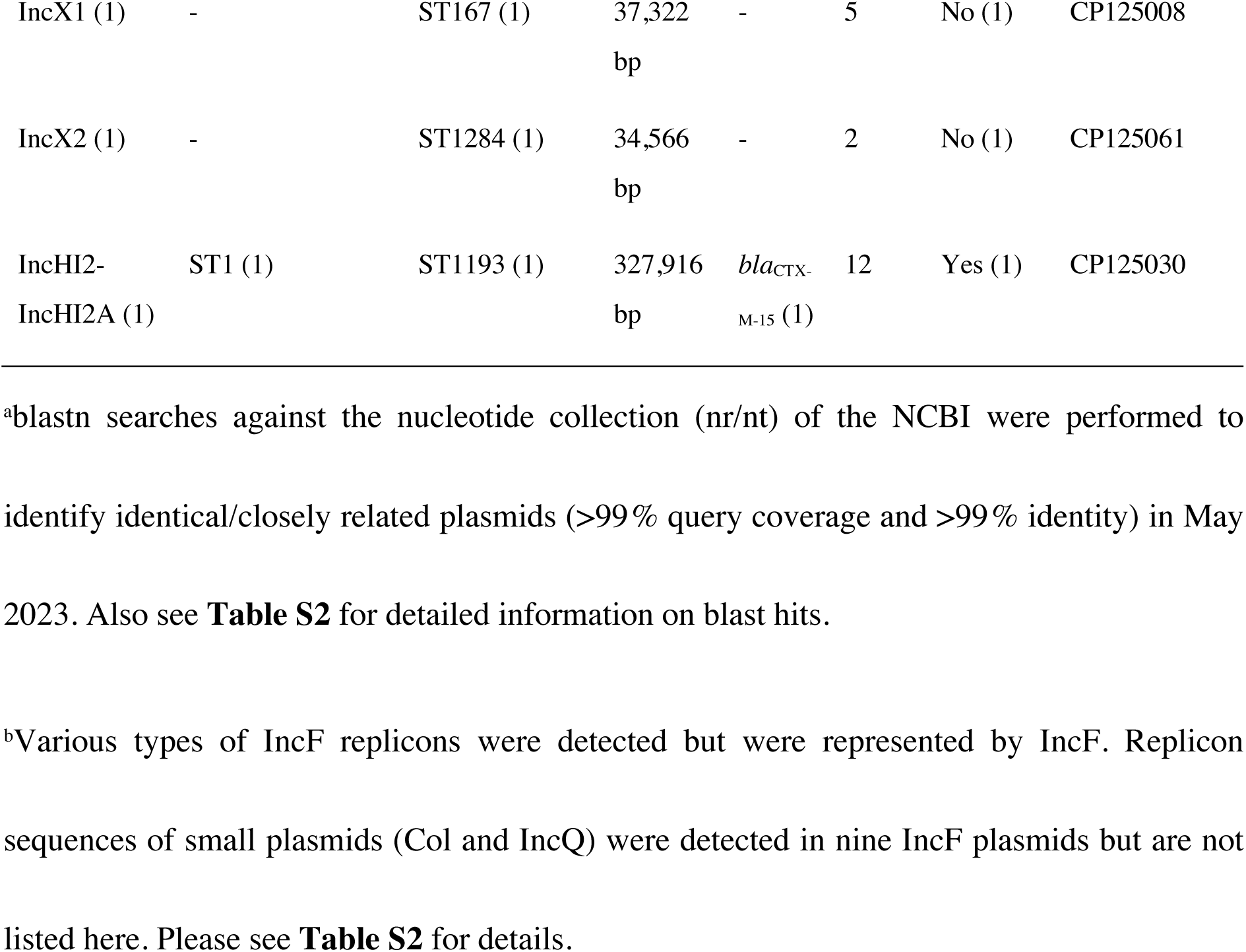
Basic characteristics of AMR plasmids in major STs.

Among the 22 completed genomes, AMR genes were detected on the chromosome in eight isolates (n = 3 ST167, n = 3 ST1193, n = 2 ST46). Notably, all eight isolates carried chromosomal *bla*_CTX-M_ (n = 7 *bla*_CTX-M-15_ and n = 1 *bla*_CTX-M-14_), and three isolates carried additional chromosomal AMR genes (**Table S2**). Detailed analyses of genetic contexts of chromosomal *bla*_CTX-M_ genes and adjacent AMR genes are provided in **Figure S3**. We note that recent studies have also shown the frequent occurrence (more than 50%) of chromosomal *bla*_CTX-M_ integration among *E. coli* isolates using both short- and long-read sequencing (Gomi et al., 2022; Milenkov et al., 2023). Chromosomal integration of AMR genes might have been previously underestimated due to the inability of PCR or short-read sequencing to determine the location of AMR genes. Long-read sequencing will become more common in the near future, which may uncover more examples of chromosomal integration of AMR genes and underline the importance of this mechanism in the stabilization and maintenance of AMR genes.

### 3.7. Study limitations

This study has some limitations. First, the clonal composition of antimicrobial-resistant *E. coli* observed in the present study might not directly reflect that of the gut flora of the underlying population due to the noise caused by e.g., differential survival of *E. coli* strains in sewer pipes/fecal sludge (Huijbers et al., 2020). Moreover, contamination from non-human animal sources in the samples cannot be excluded. Dominance of CC10 in our collection, which is prevalent not only in human feces but also in animal feces (Reid et al., 2019; Reid et al., 2017; Zingali et al., 2020), also implies this possibility. However, we think contamination from non-human animals is limited because the contribution from animal feces is lower than that from human feces in this region (Komakech et al., 2014), and *E. coli* concentrations are generally much higher in human feces than in domestic animals (Tenaillon et al., 2010). We also note that these non-human sources should also be considered in terms of the One Health concept. Inclusion of a sufficient number of fecal *E. coli* isolates from the matched human population in genomic analysis will further confirm the usefulness of fecal sludge/sewage in AMR surveillance. Second, we performed sampling over a relatively short period. Longitudinal sampling with timeframes over one year, for example, will provide information on the change of clonal composition. Third, long-read sequencing was performed only on the selected isolates from major STs. Future studies employing larger-scale long-read sequencing of isolates from various clones will elucidate AMR vectors also in minor STs. Nevertheless, our analyses revealed detailed genomic characteristics of antimicrobial-resistant *E. coli* in fecal sludge and sewage in Uganda, providing valuable insights into AMR burden in this region.

## 4. Conclusions

Here, we applied whole-genome sequencing to characterize antimicrobial-resistant *E. coli* obtained from fecal sludge and sewage in Uganda. Our study revealed that CC10, including ST167, ST10, ST1284, and ST617, were prevalent among both fecal sludge and sewage isolates, indicating this lineage significantly contributed to AMR burden in the underlying population. Global high-risk clones ST1193 and ST131 were also detected, adding potential health risks to the population. We also revealed that many of the AMR genes were carried by IncF plasmids or integrated into the chromosome by incorporating data obtained from long-read sequencing. We admit that performing whole-genome sequencing in routine surveillance is currently not realistic. However, as sequencing costs will continue to decrease, we believe that a combination of fecal sludge/sewage surveillance and whole-genome sequencing will be a powerful tool for monitoring AMR carriage in the underlying population.

## Acknowledgements

This work was supported by the Japan Society for the Promotion of Science KAKENHI (grant numbers JP18K18881 and JP19H02274). Computations were partially performed on the NIG supercomputer at ROIS National Institute of Genetics. We acknowledge the NGS core facility of the Genome Information Research Center at the Research Institute for Microbial Diseases of Osaka University for the support in DNA sequencing.

