## Supplementary Materials and methods and Supplementary Figures (S1, S2, and S3) for "Genomic surveillance of antimicrobial-resistant *Escherichia coli* in fecal sludge and sewage in Uganda"

*Corresponding authors

### Supplementary Materials and methods

#### Mash-based tree construction of clonal complex 10 (CC10) genomes

For isolates belonging to CC10, we built a tree using CC10 genomes sequenced in this study and NCBI Reference Sequence (RefSeq) CC10 genomes to gain insights into the relationships between Ugandan CC10 genomes and global CC10 genomes.

First, we downloaded allelic profiles of ST10 and related STs (max number of allele mismatches = 3) from EnteroBase to identify STs belonging to CC10. The STs marked with "ST10 Cplx" were considered as being from CC10 and saved for further analysis (n = 1,097 STs).

RefSeq *E. coli* genomes (n = 35,029) were downloaded in October 2023. Multilocus sequence typing (MLST) was performed on each genome using MLST (v2.19.0) (https://github.com/tseemann/mlst). The STs belonging to CC10, which was identified as above, were queried against ST profiles of RefSeq *E. coli* genomes to retrieve CC10 genomes. The source country information was extracted from each of the retrieved genome files, and those with valid country information were used for tree construction (n = 4,744). A tree was constructed with Mashtree (v1.0.4) (Katz et al., 2019) using the 99 CC10 genomes sequenced in the present study and the 4,744 RefSeq CC10 genomes. The tree was visualized using tvBOT (Xie et al., 2023).

#### Phylogenetic analysis of ST1193/ST131 genomes

We separately built a recombination-free tree for each of ST1193 and ST131. Isolates analyzed in other studies were also included in the trees to place our isolates in broader phylogenetic contexts (Johnson et al., 2019; Matsumura et al., 2017). Isolates from these studies were assembled and subjected to QC criteria as described in the main text, and genomes passing the QC criteria were used for tree building. For each ST, trimmed reads were mapped to a reference genome (MCJCHV-1 chromosome for ST1193 (Nielsen et al., 2018) and EC958 chromosome for ST131 (Forde et al., 2014) using snippy (v4.6.0) (https://github.com/tseemann/snippy). Recombinant regions were filtered from the snippy core-full alignment using Gubbins (v3.2.1) (Croucher et al., 2015). A maximum likelihood (ML) phylogenetic tree was built using IQ-TREE (v2.0.3) from the resulting alignment using a GTR+G+ASC model (Minh et al., 2020). The trees were visualized using iTOL (Letunic and Bork, 2021).

### Supplementary Figures

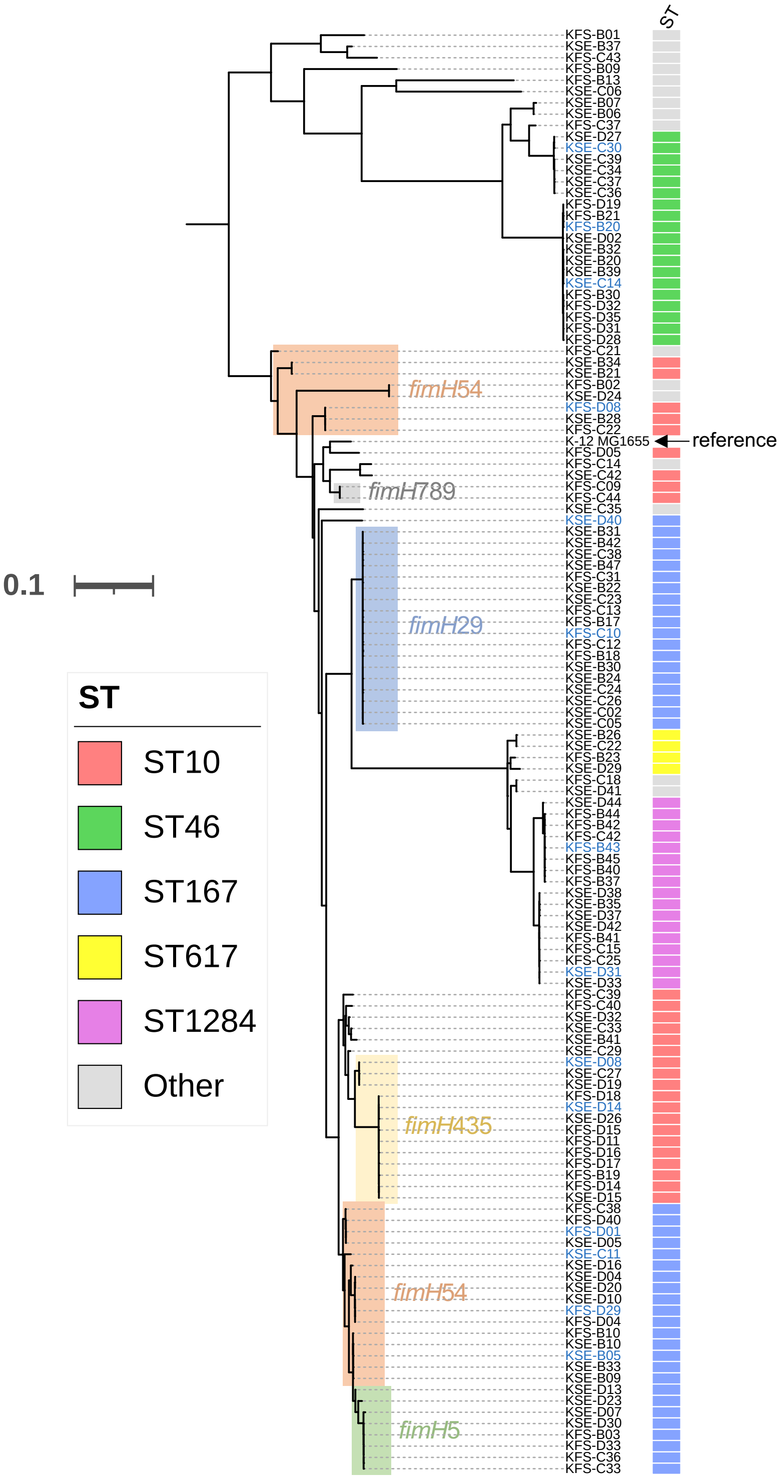


**Figure S1.** Parsnp core genome SNP tree of 127 phylogroup A isolates. K-12 substr. MG1655 (ST10, accession number: GCF_000005845.2) was used as a reference. ST10 and ST167 isolates are highlighted in the tree according to the *fimH* alleles, except for singletons. The names of isolates with completed genomes are colored blue.

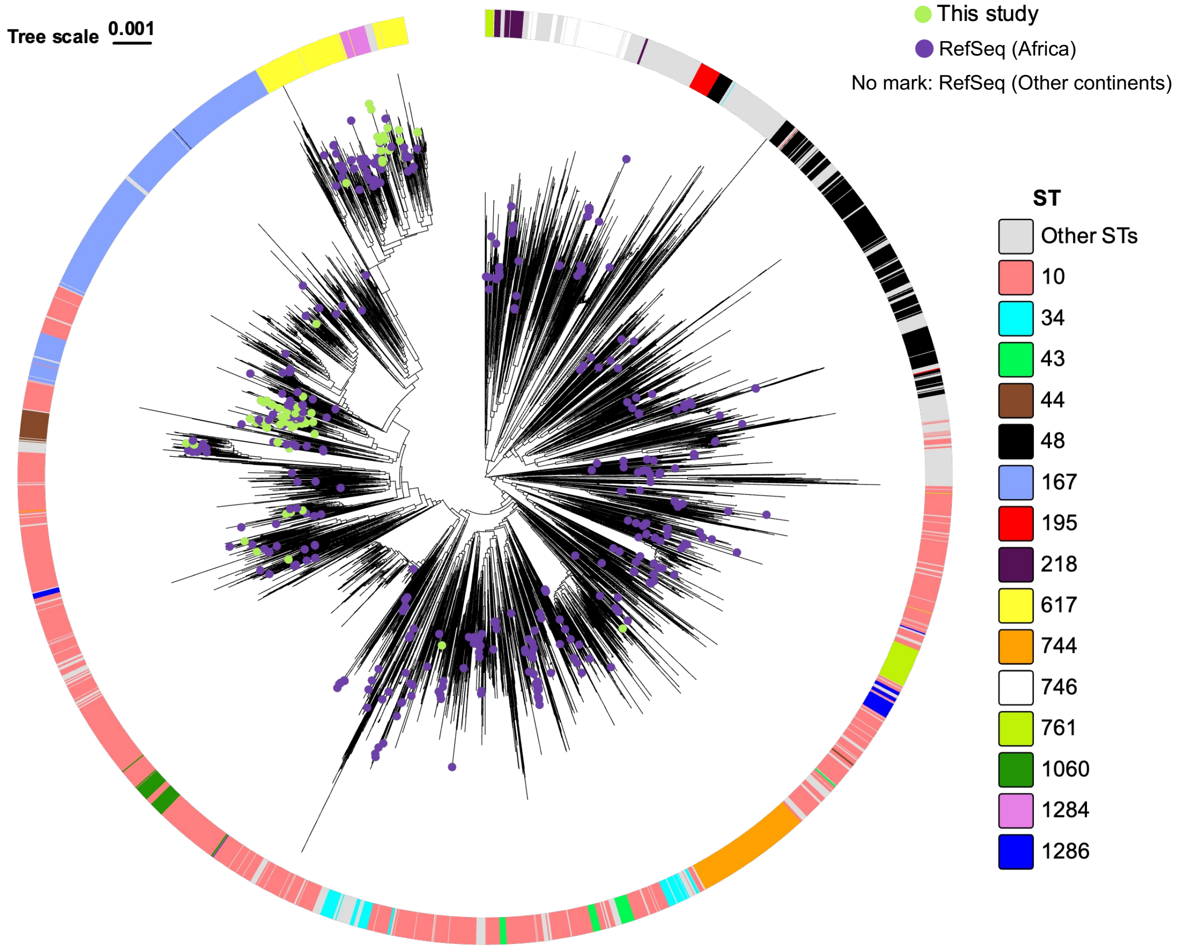


**Figure S2.** Mashtree of 4,843 CC10 genomes. The top 15 STs within CC10 are shown with different color bars and the remaining STs are shown with gray bars. The tree tips are marked with light green circles for genomes sequenced in this study and purple circles for RefSeq African genomes. RefSeq genomes from other continents were not marked to avoid complicating the figure. The tree was midpoint rooted. RefSeq CC10 genomes used in the tree construction are listed in **Table S4**.

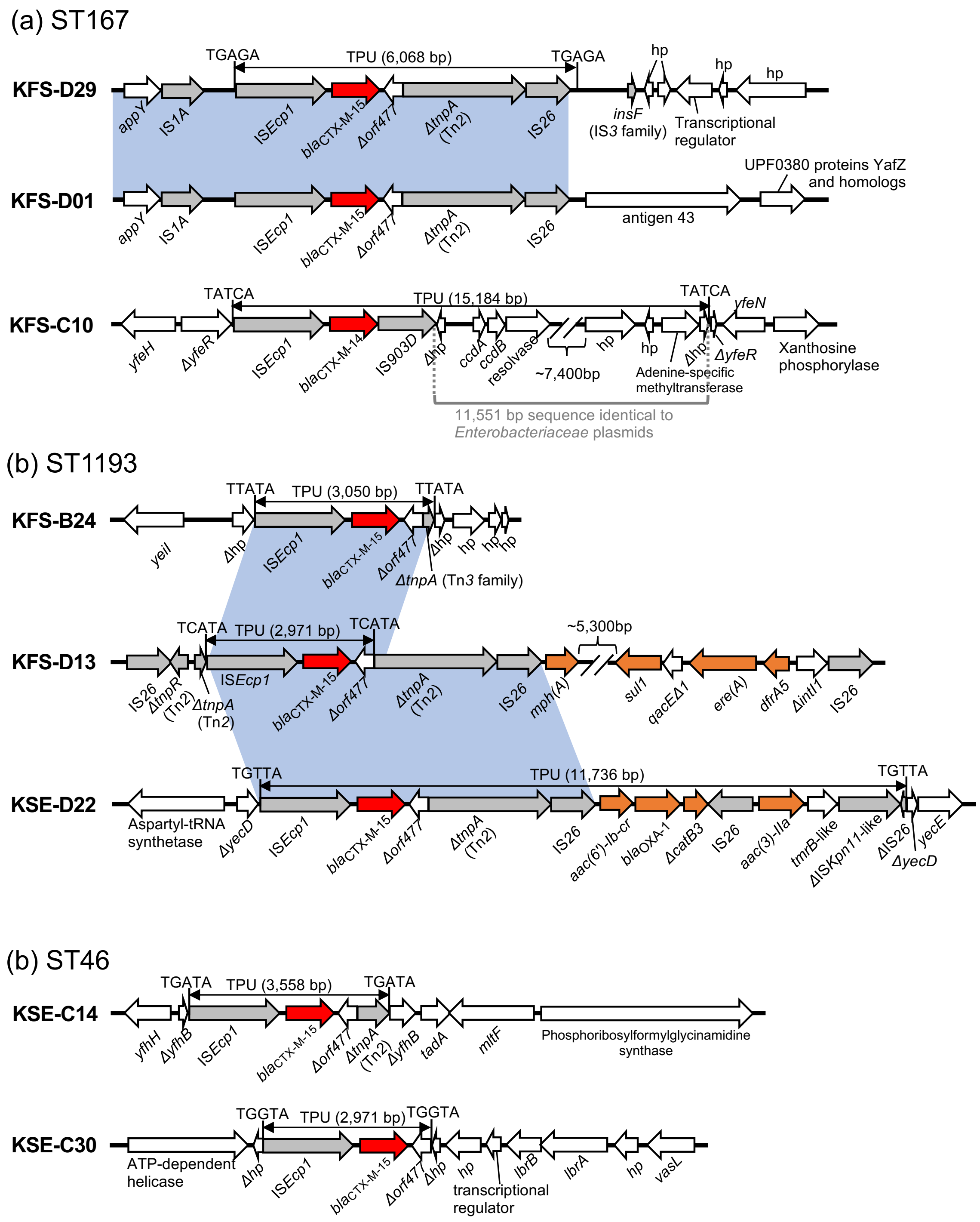


**Figure S3**. Genetic contexts of chromosomal *bla*$_{\text{CTX-M}}$ in (a) ST167, (b) ST1193, and (c) ST46. Red arrows indicate *bla*$_{\text{CTX-M}}$ genes, orange arrows indicate other AMR genes, gray arrows indicate mobile elements, and white arrows indicate other genes. The blue shaded regions indicate nucleotide sequences with 100% identity. Note that only regions related to the discussion below are shaded in blue (e.g., IS*Ecp1* in KFS-D01 and KFS-C10 are 100% identical but not shaded). The regions corresponding to transposition units (TPU) are indicated by double-headed arrows. 5-bp target site duplications (TSD) are shown next to each TPU. hp:

hypothetical protein. In KFS-D29, a 6,068 bp TPU was inserted into a sequence similar to genomic regions seen in some *Escherichia albertii* genomes, generating TSD of TGAGA. In KFS-D01, a sequence corresponding to a portion of a TPU found in KFS-D29 (from IS*Ecp1* to IS*26*) and the upstream sequence was detected, but the remaining TPU sequence and the downstream sequence were not detected. Because no TSD were detected, a TPU was not defined for KFS-D01. In KFS-C10, a TPU of 15,184 bp was inserted in *yfeR*, generating TSD of TATCA. This TPU contained an 11,551 bp sequence identical to some *Enterobacteriaceae* plasmids, suggesting incorporation of an ancestral TPU into a plasmid, followed by transposition of a new TPU (the original TPU + adjacent sequence on the plasmid) into the chromosome. In ST1193 isolates KFS-B24, KFS-D13, and KSE-D22, an IS*Ecp1*-$bla_{\text{CTX-M-15}}$-Δ*orf477*-Δ*tnpA*-IS*26* element or a portion of the element was detected. However, the TPU and TSD were different in all cases. In KFS-D13, the TPU was linked to four other AMR genes, namely *mph(A)*, *sul1*, *ere(A)*, and *dfrA5*. The region containing these four AMR genes was surrounded by IS*26*, indicating that a translocatable unit containing an IS*26* and the four AMR genes was inserted next to IS*26* which interrupts *tnpA* of Tn*2*. In KSE-D22, a 11,736 bp TPU was inserted into *yecD*, generating a TSD of TGTTA. This TPU contained $bla_{\text{CTX-M-15}}$ and four other AMR genes, *aac(6')-Ib-cr*, $bla_{\text{OXA-1}}$, Δ*catB3*, and *aac(3)-IIa*. In ST46 isolates KSE-C14 and KSE-C30, different TPU of 3,558 bp and 2,971 bp were flanked by different TSD, indicating different transposition events.
